# Ultrasound-mediated drug-free theranostics for treatment of prostate cancer

**DOI:** 10.1101/2023.09.13.555594

**Authors:** Reshani Himashika Perera, Felipe Matias Berg, Eric Chua Abenojar, Pinunta Nittayacharn, Youjoung Kim, Xinning Wang, James P. Basilion, Agata A. Exner

**Affiliations:** Department of Radiology, Case Western Reserve University, 10900 Euclid Avenue, Cleveland, OH, 44106, USA; Department of Biomedical Engineering, Case Western Reserve University, 10900 Euclid Avenue, Cleveland, OH, 44106, USA; Department of Biomedical Engineering and Department of Radiology, Case Western Reserve University, 10900 Euclid Avenue, Cleveland, OH, 44106, USA

**Author notes:** These three authors contributed equally to this work (Reshani Himashika Perera, Felipe Matias Berg, Eric Chua Abenojar).

**Keywords:** nanobubbles, PC-3 Cells, Contrast enhanced ultrasound, Prostate-specific membrane antigen (PSMA), ultrasound-mediated therapy

## Abstract

**Rationale:** Lipid-shelled nanobubbles (NBs) can be visualized and activated using noninvasive ultrasound (US) stimulation, leading to significant bioeffects. We have previously shown that active targeting of NBs to prostate-specific membrane antigen (PSMA) overexpressed in prostate cancer (PCa) enhances the cellular internalization and prolongs retention of NBs with persistent acoustic activity (∼hrs.). In this work, we hypothesized that tumor-accumulated PSMA-NBs combined with low frequency therapeutic US (TUS) will lead to selective damage and induce a therapeutic effect in PSMA-expressing tumors compared to PSMA-negative tumors.

**Methods:** PSMA-targeted NBs were formulated by following our previously established protocol. Cellular internalization of fluorescent PSMA-NBs was evaluated by confocal imaging using late endosome/lysosome staining pre- and post-TUS application. Two animal models were used to assess the technique. Mice with dual tumors (PSMA expressing and PSMA negative) received PSMA-NB injection via the tail vein followed by TUS 1 hr. post injection (termed, targeted NB therapy or TNT). Twenty-four hours after treatment mice were euthanized and tumor cell apoptosis evaluated via TUNEL staining. Mice with single tumors (either PSMA + or -) were used for survival studies. Tumor size was measured for 80 days after four consecutive TNT treatments (every 3 days). To test the approach in a larger model, immunosuppressed rabbits with orthotopic human PSMA expressing tumors received PSMA-NB injection via the tail vein followed by TUS 30 min after injection. Tumor progression was assessed via US imaging and at the end point apoptosis was measured via TUNEL staining.

**Results:** In vitro TNT studies using confocal microscopy showed that the internalized NBs and cellular compartments were disrupted after the TUS application, yet treated cells remained intact and viable. In vivo, PSMA-expressing tumors in mice receiving TNT treatment demonstrated a significantly greater extent of apoptosis (78.45 ± 9.3%, p < 0.01) compared to the other groups. TNT treatment significantly inhibited the PSMA (+) tumor growth and overall survival significantly improved (median survival time increase by 103%, p < 0.001). A significant reduction in tumor progression compared to untreated control was also seen in the rabbit model in intraprostatic (90%) and in extraprostatic lesions (94%) (p = 0.069 and 0.003, respectively).

**Conclusion:** We demonstrate for the first time the effect of PSMA-targeted nanobubble intracellular cavitation on cancer cell viability and tumor progression in two animal models. Data demonstrate that the targeted nanobubble therapy (TNT) approach relies primarily on mechanical disruption of intracellular vesicles and the resulting bioeffects appear to be more specific to target cancer cells expressing the PSMA receptor. The effect, while not lethal *in vitro*, resulted in significant tumor apoptosis *in vivo* in both a mouse and a rabbit model of PCa. While the mechanism of action of these effects is yet unclear, it is likely related to a locally-induced immune response, opening the door to future investigations in this area.

## Introduction

Prostate cancer (PCa) is the second most common solid malignancy in men globally and the second highest contributor to the mortality rate in Western countries[1–3]. Although localized PCa is not lethal, it causes a large spectrum of aggressive diseases which contribute to men’s mortality [4]. Early detection of PCa is limited by nonspecific screening tests, such as prostate-specific membrane antigen (PSA) testing and trans-rectal ultrasound-guided (TRUS) biopsy which often produce false positive and false negative results, respectively [5]. Nonetheless, these conventional PCa screening methods are essential to improving accurate early-stage PCa diagnosis and reducing mortality. High-risk PCa (Gleason score ≥8, PSA > 20 ng/ml or clinical stage ≥T3a) has more than 35% cumulative mortality at 15 years [6,7]. Thus, establishing an effective early diagnosis method along with a suitable treatment regimen is beneficial for patients with high-risk, localized PCa.

Standard therapies for aggressive PCa include surgical resection, stereotactic body radiotherapy (SBRT), androgen deprivation therapy (ADT), and combined therapy, depending on the risk level of the patient. Apart from localized therapy, systemic therapy such as hormonal therapy, chemotherapy, and immunotherapy are other approaches in PCa treatment [8,9]. Despite recent successes, there remain significant limitations to these strategies due to side effects, complex logistics, and treatment costs. Radiation therapy and surgical resection are associated with side effects such as urinary dysfunction, erectile dysfunction, and damage to surrounding tissues [10]. Therefore, less disruptive options with minimum side effects are needed to improve treatment efficacy and quality of life following treatment of PCa [11].

Previously, we reported a contrast-enhanced ultrasound (CEUS) imaging technique to improve the imaging capabilities of PCa using nano-sized prostate-specific membrane antigen (PSMA)-targeted ultrasound (US) contrast agents [12,13]. PSMA is a membrane-bound, type II integral protein that is highly overexpressed in PCa compared to normal prostate tissue [14–16]. Prior work has demonstrated that PSMA-targeted nanobubbles (PSMA-NBs) selectively accumulate and internalize into cancer cells of the PSMA-expressing tumor [12,13,17]. When placed in an acoustic field, nanobubbles can oscillate (stable cavitation) and / or collapse (inertial cavitation) depending on the incident ultrasonic energy. In diagnostic imaging, bubbles typically oscillate, but with sufficiently high driving amplitude, can violently collapse. Bubble collapse generates a transient high-pressure jetstream capable of puncturing through neighboring cells [18].

Acoustic cavitation has been cited as the most important non-thermal ultrasound mechanism that occurs through the generation of local acoustic streaming, which changes the biological system [19–23]. This triggered bubble destruction has been widely used to transport drugs and genes to the target for increased therapeutic efficacy[24,25]. However, US-triggered inertial cavitation is typically non-specific to the disease lesion, and focused ultrasound equipment is required to localize the cavitation to a target site. In contrast, specific targeting of nanobubbles to a biomarker such as PSMA expressed solely on the target cells, can result in disease-specific agent localization without the need for focused ultrasound. Here, when unfocused therapeutic ultrasound (TUS) is applied after targeted NBs bind and/or internalize into the target cells, NBs can be cavitated, resulting is selective cell elimination.

In this work we combine the receptor-mediated endocytosis of PSMA-NBs with externally applied US as an effective treatment strategy for PCa. We evaluate the strategy *in vitro, in vivo* in a mouse model of human PCa expressing PSMA, and in an orthotopic PCa model expressing PSMA in rabbits [26]. We hypothesize that the cavitation of internalized PSMA-NBs will selectively damage the tumor cells and contribute to better treatment efficacy with minimum side effects.

## Methods

### Preparation and characterization of contrast agents

The preparation and characterization of NBs have been reported previously [27–29]. Briefly, a cocktail of lipids including DBPC (Avanti Polar Lipids Inc., Pelham, AL), DPPE, DPPA (Corden Pharma, Switzerland), and mPEG-DSPE2000 (Laysan Lipids, Arab, AL) were dissolved in propylene glycol (PG, Sigma Aldrich, Milwaukee, WI), glycerol and PBS and sealed in a vial. Then the gas from the sealed vial was exchanged with C_3_F_8_ (Electronic Fluorocarbons, LLC, PA), and the vial was subjected to mechanical agitation (Vialmix^®^). NBs were isolated by differential centrifugation. PSMA-NBs were formulated by incorporating DSPE-PEG-PSMA-1 into the lipid cocktail mixture [13]. PSMA-NBs and NBs were characterized with resonant mass measurement (RMM; Archimedes^®^, Malvern Panalytical) and validated using HPLC and Matrix-assisted laser desorption/ionization time-of-flight mass spectroscopy (MALDI-TOF-MS) as previously described [13,27]. Rhodamine-labeled NBs were prepared by mixing DSPE-Rhodamine (50 μL) into the lipid solution.

### Cell culture

Retrovirally-transformed PSMA-positive (PC3pip) cells and transfection-control PSMA-negative (PC3flu) cells were originally obtained from Dr. Michel Sadelain (Memorial-Sloan Kettering Cancer Center, New York, NY). The two cell lines were checked and authenticated by Western blot. Cells were grown at 37°C and 5% CO_2_ and maintained in a complete RPMI1640 medium (Invitrogen Life Technology, Grand Island, NY).

### Confocal imaging of internalized PSMA-NB in PSMA positive cells after TUS

PSMA+ and PSMA-cells were seeded in glass-bottom petri dishes (MetTek Corporation, Ashland, MA, USA) at a density of 5×10^4^ cells/well. After 24 hrs, Rhodamine-tagged PSMA-NBs (40,000 NBs per cell) were added to the cells and allowed to incubate for 1 hour. Following incubation, cells were washed with PBS for 3 times. To visualize late endosomes and lysosomes, RPMI with 5% fetal bovine serum and 5 μM Lysotracker deep red (ThermoFisher Scientific) was used following the manufacturer’s instructions. TUS (Mettler Sonicator 740 Therapeutic Ultrasound, San Diego, CA) was applied for 10 min from the top of the petri dish (covered with parafilm in full contact with the media, and using coupling gel) with the parameters of 3 MHz, 2.2 W/cm2, and 10% duty cycle (DC). After the treatment, cells were fixed with 4% paraformaldehyde for 10 min. Cells were washed with PBS and stained with DAPI mounting medium (Vector Laboratories, Burlingame, CA) and observed using a confocal microscope (Leica DMI 4000B, Wetzlar, Germany) equipped with the appropriate filter sets (DAPI, Rhodamine, and Lysotracker channels).

### WST-1 cell viability assay

PSMA+ and PSMA-cells were seeded in a 96-well plate (200 μl of cell suspension of 1×10^5^ cells/ ml density). After PSMA-NB co-incubation with cells for 1 hr, TUS treatment was applied (with the same parameters as above) from the top of the plate. The cell viability was measured at 24 hrs and 48 hrs post-treatment. Cells were incubated with WST-1 reagent according to the manufacturer’s protocol. The absorbance was measured at 450 nm wavelength.

### Subcutaneous tumor implant in mouse model

Mice were handled according to the protocol approved by the Institutional Animal Care and Use Committee (IACUC) at Case Western Reserve University and were in accordance with all applicable protocols and guidelines for animal use. Male athymic nude mice (4-6 weeks old) were anesthetized with inhalation of 3% isoflurane with 1 L/ min oxygen and were implanted subcutaneously with 1×10^6^ of PSMA+ and PSMA-cells in 100 µL matrigel. For the non-survival, immunohistochemical analysis and for US imaging (n=8), PSMA+ and PSMA-tumors were inoculated in both flank areas (left and right) of the same mouse (dual tumor model). For survival studies, PSMA+ and PSMA-tumors were inoculated in separate animals (single tumor model). Animals were observed twice per week.

### *In vivo* tumor imaging in mouse model

Mice were inoculated with PSMA+ and PSMA-tumors in the flank area (dual tumor model, Figure 1). The animals were imaged when the tumor diameter was ∼8 mm (n=2). The US probe (PLT-1204BT, AplioXG SSA-790A, Toshiba Clinical Medical-Imaging Systems, Otawara-Shi, Japan) was placed to visualize the PSMA+ and PSMA-tumors in the same field of view. Undiluted PSMA-NBs (200 μl) at 4×10^11^ NBs/mL were administered via tail-vein injection. After injection, the tumors were imaged using contrast harmonic imaging (CHI, frequency: 12.0 MHz; MI: 0.1; dynamic range: 65dB; gain: 70dB; frame rate: 0.2 fps) for 3 min to confirm NB injection and NB influx into the tumor. PSMA-NBs were then left to circulate freely and accumulate in the tumor for 1hr. without scanning, but with the probe at the same position. After 1hr, tumors were scanned again to examine the accumulation of bubbles in both PSMA+ and PSMA-tumors. The tumor areas were delineated by drawing regions of interest (ROIs) and the average signal intensity of the drawn ROIs was obtained.

**Figure 1.**
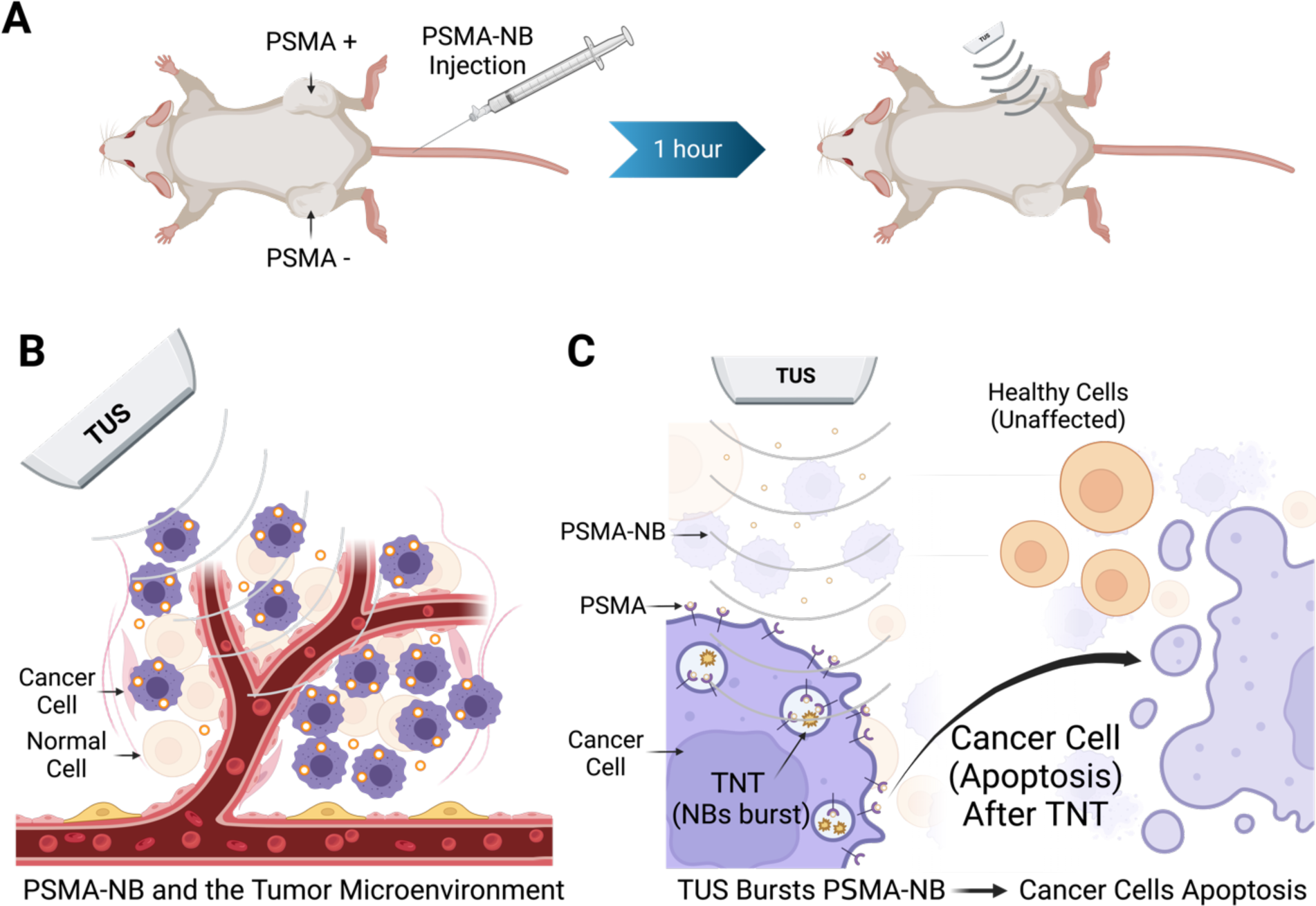
Schematic diagram showing the experimental approach for targeted NB therapy (TNT) with PSMA-NB injection (left) followed by TUS irradiation (right) (A). PSMA-targeted NBs selectively internalized into PCa cells. When circulating NBs wash out after 1 hour, the remaining NBs in tumors are cavitated using low frequency therapeutic ultrasound (B). Intracellular NB cavitation induces effects which ultimately result in tumor cell apoptosis and reduction of tumor progression. (C).

### Tumor treatment in mouse model

Animals were inoculated with PSMA+ and PSMA-tumors (dual tumor model) in the flank area on both sides (Figure 1). Two weeks after inoculation, animals received either 200 μL of undiluted PSMA-NB or PBS via the tail vein. 1 hr. post-injection, the PSMA positive TNT group (PSMA-NB + TUS treatment, n=3) received 10 min of therapeutic US (TUS) in only the PSMA positive tumor with the same parameters as above (3 MHz, 2.2 W/cm^2^, 10% duty cycle (DC), 5cm^2^ probe area). The PSMA negative tumor on the same animal was considered as the PSMA-NB-only treatment (no TUS treatment). Similarly, for the PSMA negative-TNT group (n=3), TUS was applied 1 hr. post-injection to the PSMA-tumor while the PSMA+ tumor was treated with only PSMA-NBs. For animals injected with PBS, both tumors were treated with TUS and considered as the TUS-only treated tumors (n=2). The TUS treatment order for PSMA positive and PSMA negative was switched for control animals.

### Immunohistochemistry and histological evaluation in mouse model

Tumors were harvested 24 hr. after the treatment, fixed in 4% paraformaldehyde, and embedded in optimal-cutting-temperature compound (OCT Sakura Finetek USA Inc., Torrance, CA). Tissues were then cut into 12 μm thick slices and washed (3X) with PBS. To evaluate apoptotic cell death, a TdT-mediated dUTP nick-End Labeling (TUNEL) assay was used following the manufacturer protocol (Abcam, Boston, MA). The apoptotic or TUNEL-positive cells/areas in the tumor that were stained with dark brown color were counted using ImageJ. The ratio of TUNEL-positive cells/areas relative to the whole tumor area was reported as a percentage. H&E staining was performed following the standard protocol and the whole-tumor tissue slides were scanned at 20x objective with the Axioscan Z1 slide scanner (Zeiss Inc., Oberkochen, Germany) under the same exposure times.

### *In vivo* survival assessment in mouse model

Survival studies were performed to examine the TNT treatment efficacy for PCa cancer survival. For the survival studies, either PSMA+ tumor or PSMA-tumor was inoculated in the flank area in separate mice (single tumor model). PSMA-positive tumor-bearing mice and PSMA-negative tumor-bearing mice were treated with either PSMA-NB plus therapeutic ultrasound (designated as TNT), PSMA-NB alone (without ultrasound) or with therapeutic ultrasound only (designated as TUS only). Two weeks after inoculation, animals were treated with either TNT, TUS, or no treatment. All mice underwent the same protocol for the anesthetization (as explained above), injection of PSMA-NB, and the TUS treatment. The TUS parameters were 3 MHz, 2.2 W/cm2, and 10% duty cycle (DC) for 10 min, consistent with methods above. The same treatments were performed for each animal 3 more times at intervals of 3 days from the starting date of the treatment. The tumor volume was measured from US images 2-3 per week. Animals were euthanized and tumors were collected before the tumor volume exceeded the IACUC standards (when the tumor size reaches > 15 mm diameter).

### Orthotopic tumor implant in rabbit model

A total of 14 sexually mature male White New Zealand rabbits (Charles River Laboratories, Garfield Heights, OH, United States) were included in this study, divided into two groups: the treatment group (n=7) and the control group (n=7). All animal procedures were performed under inhalatory anesthesia (isoflurane 2%) unless otherwise noted. The tumor inoculation and immune suppression procedures were performed as previously described [26]. In brief, all animals were subjected to immunosuppression with cyclosporine (10 mg/kg) administered subcutaneously. The immunosuppressive regimen was initiated one day before the tumor inoculation and was continued with daily injections of cyclosporine until the end of the protocol. PSMA+ cells expressing green-fluorescent protein (GFP) were inoculated via a transabdominal approach under US guidance with 21-gauge needles (50.8 mm in length). The needles were primed with cells before the beginning of the procedure. Once the location of the needle tip was confirmed to be inside the prostate gland by the US, 100 μL (8 x 10^6^ cells) were injected.

### Tumor imaging in rabbit model

For the rabbit model, all US images were acquired using a Siemens Acuson S3000 (Siemens Healthineers, Erlangen, Germany) and a 6-18 MHz linear array probe (18L6 Probe). Prior to imaging, the abdominal hair of the animals was trimmed and removed using hair removal cream (Church & Dwight, Ewing Township, NJ, United States). The prostate gland was located in the lower abdomen/pelvis of the animal, situated caudally to the bladder and posteriorly to the vas deferens ampulla and urethra (Figure 5B). After locating the prostate gland, 0.8 mL/kg of PSMA-NB (4.0 x 10^11^ NBs/mL) was injected using a peripheral vein catheter placed in the animal’s ear. The prostate region was continuously imaged for 2 min (frequency: 8 MHz, MI: 0.10, 1 fps) to evaluate the contrast wash-in. After 2 min, the imaging was interrupted to let NBs circulate and accumulate for 30 min. Weekly US imaging was performed for tumor monitoring. B-mode, color Doppler, and contrast-enhanced ultrasound (CEUS) images were acquired.

### Treatment of orthotopic tumors in rabbit model

After the CEUS (as explained above) imaging was concluded, the therapeutic ultrasound probe was positioned over the prostate gland area, previously marked based on US images. For the treatment group (n = 7), treatment was initiated 30 min after the PSMA-NB injection to allow for NBs cell incorporation. The treatment was performed for 15 min using the following TUS parameters: frequency 1 MHz, 2.2 W/cm2, 10% duty cycle (DC). The therapeutic US probe measured 4.2 cm in diameter (10 cm^2^ area) and had a nominal depth penetrance of 5 cm. For the control group (n = 7), animals were injected with PSMA-NB alone (no TUS).

### Fluorescence imaging and histology for rabbit model

Animals were euthanized with pentobarbital overdose and the prostate, proprostate, and paraprostate were harvested *en bloc* with the bladder and urethra. Tissues were washed with PBS and imaged with the Maestro Macro-Fluorescence system. After fluorescence imaging, tissues were fixed in 10% formalin for 24 hrs. and processed using standard protocols. Tumor volume was calculated using US images. The PSMA expression was assessed by immunohistochemistry using a primary rabbit polyclonal antibody anti-PSMA and a secondary antibody conjugated to horseradish peroxidase. Fluorescent images obtained with the Maestro system were analyzed using the Maestro software, and GFP signal intensity was quantified for tumor location validation. H&E staining was performed as explained above.

### Statistical Analysis

Graphs generated using GraphPad PRISM^®^. Statistical analysis was performed with R Software (version 4.3). Unpaired Student’s t-test (two-tailed), Welch’s Test were used to compare 2 groups with normal variances and not normal variances, respectively. ANOVA single factor was used to compare 3 groups with unless otherwise noted. Data are presented as a mean ± SEM (standard error mean) unless otherwise indicated. The experiments were repeated three times unless stated otherwise.

## Results

### Nanobubble preparation and characterization

Nanobubbles were prepared and functionalized with the PSMA-1 ligand and characterization of the lipid-ligand conjugation was carried out as previously reported [13]. The diameter of PSMA-NBs was 277 ± 97 nm as characterized by resonant mass measurement (RMM). The standard deviation (SD) of NB diameter was calculated from the mean bubble diameter for three formulations. The concentration of PSMA-NB was 4 x 10^11^ ± 2.45 ×10^10^ NB/ ml. Validation of the RMM analysis and its optimization for use in NB characterization has been previously described [13,28]. PSMA-NB accumulation in the tumor, internalization into cancer cells, and bubble cavitation with the application of TUS are illustrated in Figure 1.

### Confocal imaging of cellular internalized PSMA-NB and after TUS and treatment evaluation *in vitro*

The localization of PSMA-NBs in the cellular compartments and the associated change in cellular structures after therapeutic US were evaluated using confocal microscopic imaging (Figure 2). Our previous fluorescence imaging data showed that the PSMA-NBs are selectively internalized by the PSMA+ cells compared to the plain NB [13]. Similarly, the confocal imaging results presented here show high internalization and localization to cellular compartments for PSMA-NB co-incubated PSMA+ cells compared to PSMA-cells. (Figure 2A, supplementary Figure S1 and S2, 100X). The fluorescence dye, Lysotracker^®^, was used to stain late endosomes and lysosomal structures, and is pseudo-colored in green in the images to avoid overlap with the Rhodamine-labeled NBs (shown in red). Co-localization (yellow) of late endosomes/lysosomes (green) and PSMA-NBs (red) was observed in PSMA-expressing cells. PSMA negative cells showed non-specific uptake of PSMA-NB with lesser late endosomal/lysosomal co-localization. Twenty-four hours post incubation, PSMA+ cells still showed a higher amount of co-localized PSMA-NBs (yellow staining) compared to the PSMA-cells, indicating the presence of PSMA-NB inside the cellular compartment (Figure 2C). Prior work measured entrapped C3F8 gas in the PSMA+ cells, suggesting the nanobubbles are still active following internalization into cells. [17] Furthermore, as shown in Figure 2B, after the PSMA-NB incubation and TUS treatment, the interruption of the cellular compartment was visible in PSMA+ cells. Confocal imaging results confirmed the internalization of NBs and the disruption of cellular compartments after TUS application, indicating cellular damage resulting from NB cavitation (concept demonstrated in Figure 1). Furthermore, 24 hrs post-TNT treatment (PSMA-NB + TUS), PSMA+ cells showed disrupted cellular compartments with more PSMA-NB signal (red color) in most of the cells compared to the PSMA+ cells at 24 hr. post PSMA-NB incubation (Figure 2D). In contrast, small amounts of PSMA-NB signal and cellular compartments were visualized in PSMA-cells after 24 hrs post incubation with PSMA-NBs and post TNT treatment. Cell viability decreased slightly in PSMA+ cells treated with TNT (reduction of 24.6 ± 1.4% 48 hrs after treatment). Cell viability following TNT treatment in control groups was not affected (supplementary Figure S3).

**Figure 2.**
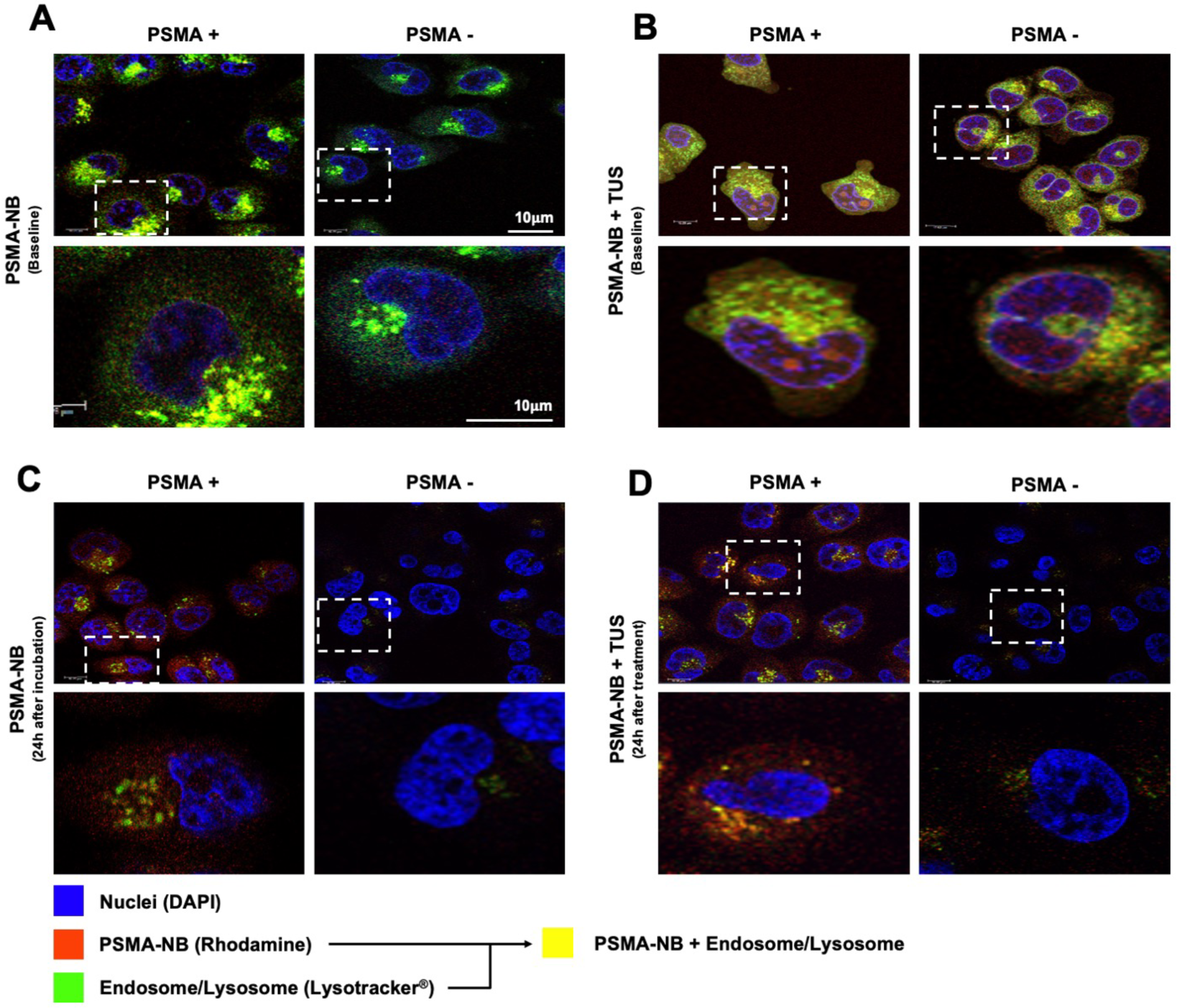
Confocal images of PSMA-NB distribution in PSMA+ cells and PSMA-cells after in vitro co-incubation with fluorescent PSMA-NB (immediately after incubation) (A), after co-incubation with PSMA-NB and therapeutic US (TUS) application (10 min, 2.2 power, 3 MHz, and 10% duty cycle) (B), PSMA-NB distribution after 24 h (C) and 24 h post incubation and TUS treatment (D); (blue-nuclei, red-NB, green-late endosome/lysosomes, and yellow-co-localized PSMA-NBs with endosome/lysosomes). 100x Higher amplification for each image in the 1^st^ and 3^rd^ is provided on the 2^nd^ and 4^th^ rows (respectively).

### *In vivo* ultrasound imaging showing PSMA-NB accumulation after 1 hr. in mouse model

*In vivo,* CEUS experiments were performed to evaluate the accumulation of PSMA-NBs in both PSMA+ and PSMA-tumors at 1 hr post-injection. Rapid contrast enhancement was observed within both tumors and peak accumulation of agents in the tumors was observed at 3 min (Figure S4). After 3 min, the US imaging was paused to allow NBs to freely circulate without US exposure. At 1 hr. post-injection, imaging acquisition was restarted to visualize the contrast accumulation in the tumors. 25% higher enhancement was observed in the PSMA+ tumor compared to the PSMA-tumor suggesting greater NB accumulation and retention in the PSMA+ tumors. (Figure S4).

### TUNEL and H&E staining in mice

To visualize apoptosis in tissue sections, TUNEL staining was conducted 24 h after TNT treatment. TUNEL assay stains apoptotic cell nuclei dark brown. As shown in Figure 3 (A1-A6, column 1), we observed significantly higher numbers of dark brown nuclei in TNT-treated PSMA positive tumor tissues compared to TNT-treated PSMA negative tumor tissues (78.52 ± 9.3% Vs 30.48 ± 6.012%, p < 0.01), indicating a marked increase in apoptosis. The TNT-treated PSMA+ tumor had a high percentage of apoptosis compared to all other groups tested. Control treatments demonstrated that TUS alone or PSMA-NB alone did not stimulate apoptosis.

**Figure 3.**
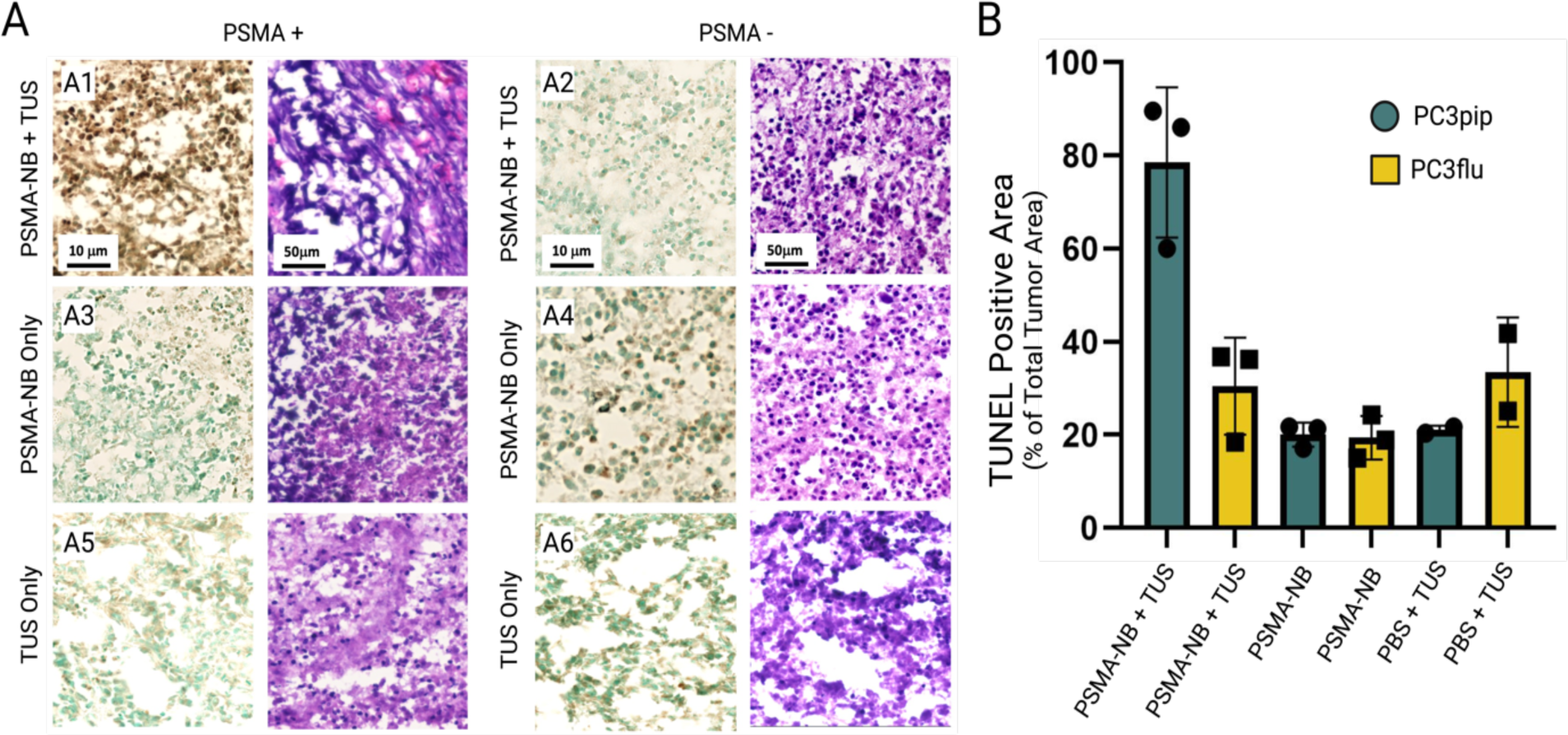
TUNEL and H&E stained images of tumors treated with TNT, PSMA-NB or TUS alone (A). The percent of TUNEL positive area compared to the whole tumor area (B). Note only TNT with PSMA+ tumors generate significant apoptosis.

H&E staining was performed to examine the comprehensive histological characteristics of PSMA+ and PSMA-tumor xenografts after the TNT, PSMA-NB, and TUS treatments (Figure 3). As shown in Figure 3 (A1-A6, column 2), NB-only and TUS-only treated PSMA+ and PSMA-tumors showed clear intact cell nuclei. PSMA+ tumors treated with TNT exhibited nuclear fragments and malformed nuclei (Figure 3A1, column 2).

### *In vivo* survival in mice

PSMA-NBs combined with unfocused, low frequency TUS were used to demonstrate the efficacy of targeted intracellular NB cavitation treatment (TNT) *in vivo* in an immunocompromised athymic mouse model. The timeline of the treatment is shown in Figure 4A. Figure 4B1-4B2 show the long-term (80 days) changes in tumor progression and animal survival of mice with PSMA expressing tumors compared to the untreated control and TUS-only treatment without injection of NBs. Control tumors grew rapidly and reached the planned cutoff size (endpoint; tumor diameter ∼15 mm) 4 weeks after tumor inoculation. Progression of PSMA+ tumors treated with TNT-was significantly slower than both the control and TUS-only treated groups (p < 0.001). After 38 days, mice in the TUS-only treated control group reached the tumor volume endpoint, while the tumor growth was diminished in the TNT-treated group, where 30% of animals survived for more than 60 days. B-mode ultrasound images of tumors were also performed throughout the course of the study (Figure S5). From B-mode images, progressive growth of necrotic areas with tumor progression is apparent. This is most evident in the control group and is consistent with the tumor’s rapid growth and central necrosis. Twenty percent of TNT-treated PSMA+ tumor animals showed reduced tumor growth up to 75-days post treatment (Figure S5B).

**Figure 4.**
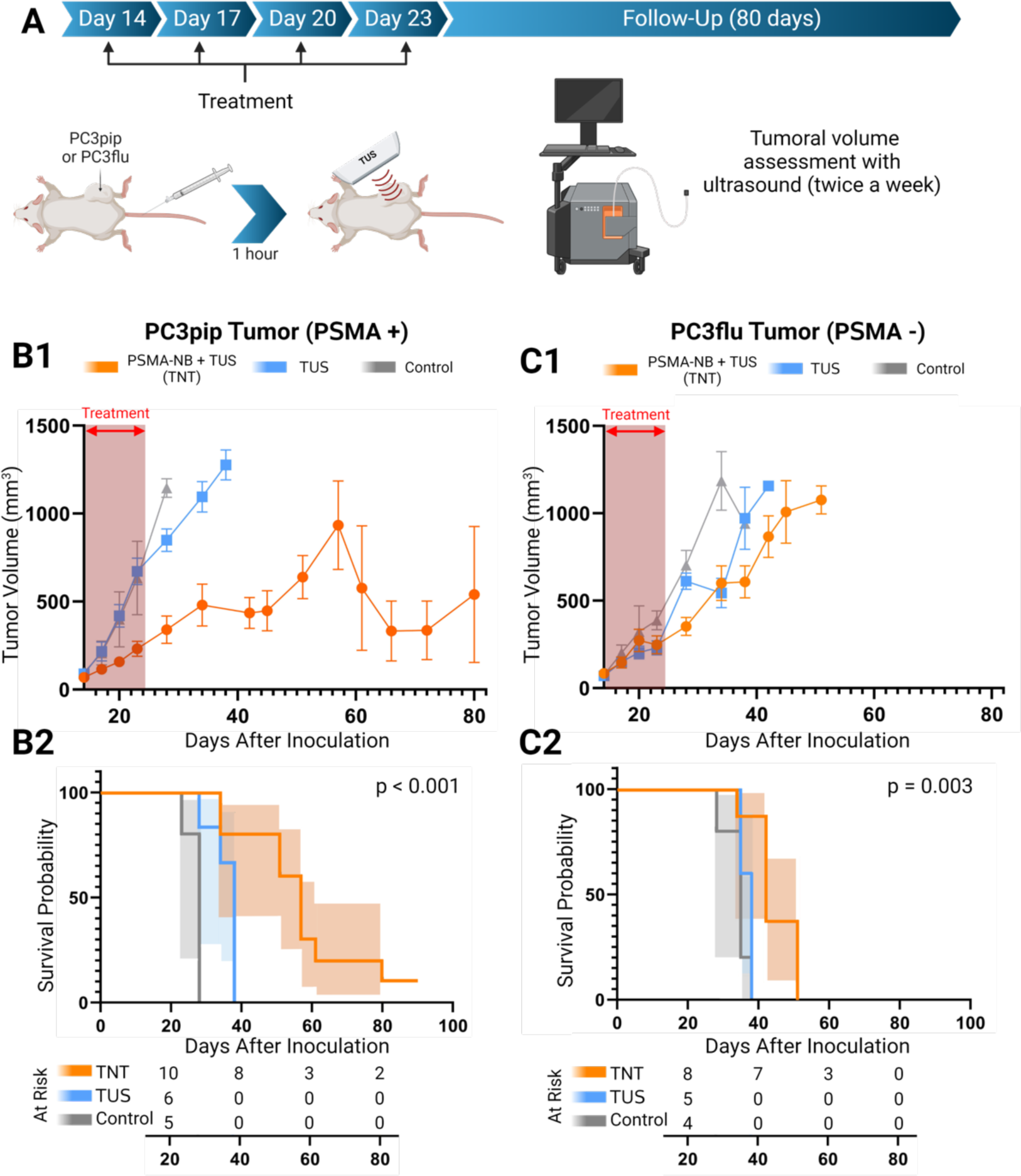
Schematic diagram of the experimental setup and timeline (A). Tumor growth curve (mean ± SEM) of PSMA+ tumor mice (PSMA+ TNT (n=10), PSMA+ TUS (n=6), PSMA+ control (n=5)) (B1) and PSMA-tumor mice (PSMA-TNT (n=8), PSMA-TUS (n=5), PSMA-control (n=5)) (C1) after treatment. Survival curve with 95% C.I. (shaded area) of PSMA+ tumor (B2) and PSMA-tumor-bearing mice (C2) after TNT treatment. Red dashed arrows indicate the days treatment was performed. P-values for log-rank test. Bothe TNT and TUS-Only treatments were applied every 3 days starting from 14^th^ day to 24^th^ day (four treatments) after tumor inoculation.

For further analysis, tumors were divided into two groups depending on the initial tumor size (first treatment day): large tumors (> 60mm^3^) and small tumors (< 60 mm^3^ ). As shown in Figure S6A and S6C, mice with smaller tumors (red) showed a better response to the TNT treatment compared to mice with larger tumors (blue). In the small tumor group, 65% of animals showed hindered tumor growth after the TNT treatment. The TNT-treated group of PSMA+ tumors demonstrated significant tumor volume reduction at 28-day post-inoculation compared to both TUS-only and control groups (p < 0.001). The Kaplan-Meier survival analysis showed consistent prolonged survival of the PSMA+ tumor mice that received combined treatment compared to the control and TUS-only treated group.

In contrast to PSMA-expressing tumors, PSMA-negative tumors treated with TNT exhibited faster growth rates. However, the tumors did show a significant initial decrease in tumor progression compared to the control groups treated with therapeutic ultrasound (TUS) alone and the control groups of the same tumor type (Figure 4C1, C2, and Figure S6B, S7A), At 35 days post-treatment and later, no significant difference in tumor growth was observed among the treated PSMA-tumors and corresponding controls (Figures S6B and S6D). Additionally, B-mode images (Figure S7) revealed rapid tumor growth with increasing necrotic areas in the tumor. Notably, Kaplan-Meier survival analysis demonstrated an improvement in survival for the TNT-treated PSMA-tumors (50 days after inoculation) compared to the other groups (p = 0.003). Conversely, the survival of TUS-treated groups without NBs, both for PSMA-positive (PSMA+) and PSMA-tumor-bearing animals, was similar at the 38-day endpoint. However, upon discontinuation of the treatment, PSMA negative tumors exhibited rapid regrowth (Figure S6B).

### *In vivo* ultrasound imaging and treatment for rabbit model

Tumors expressing human PSMA were inoculated inside the rabbit prostate gland, followed by weekly irradiation of the prostate using targeted ultrasound (TUS) following priming with PSMA-NBs (Figure 5). Following inoculation, tumors grew both within the prostate gland and also outside of the gland, likely due to seeding of cells along the needle track. Remarkably the treatment significantly inhibited tumoral growth for both intraprostatic and extraprostatic/intraperitoneal tumor locations. In the treatment group, rabbits with intraprostatic tumors showed a 90% reduction in tumor size compared to the control group at the study endpoint of 5 weeks (p = 0.090, mean difference ± SEM: 260 ± 130 mm3) (Figure 6A). Notably, the control group exhibited a higher risk of developing intraprostatic tumors (RR: 1.67; 95% C.I.: 0.62 - 4.42). The effects on extraprostatic tumors were even more pronounced, with tumors in the treatment group being 94% smaller compared to the control group (p = 0.003, mean difference ± SEM: 1440 ± 468 mm3) (Figure 6B). In the control group, all animals developed extraprostatic tumors, while only 28.6% of animals in the treatment group developed extraprostatic tumors (RR: 3.50; 95% C.I.: 1.08 - 11.29). MRI and ex vivo fluorescence imaging were used to confirm the tumor localization of PSMA-NBs within the treatment area (Figure S8). Supplementary material (Figure S9) provides individual growth curves for each tumor.

**Figure 5.**
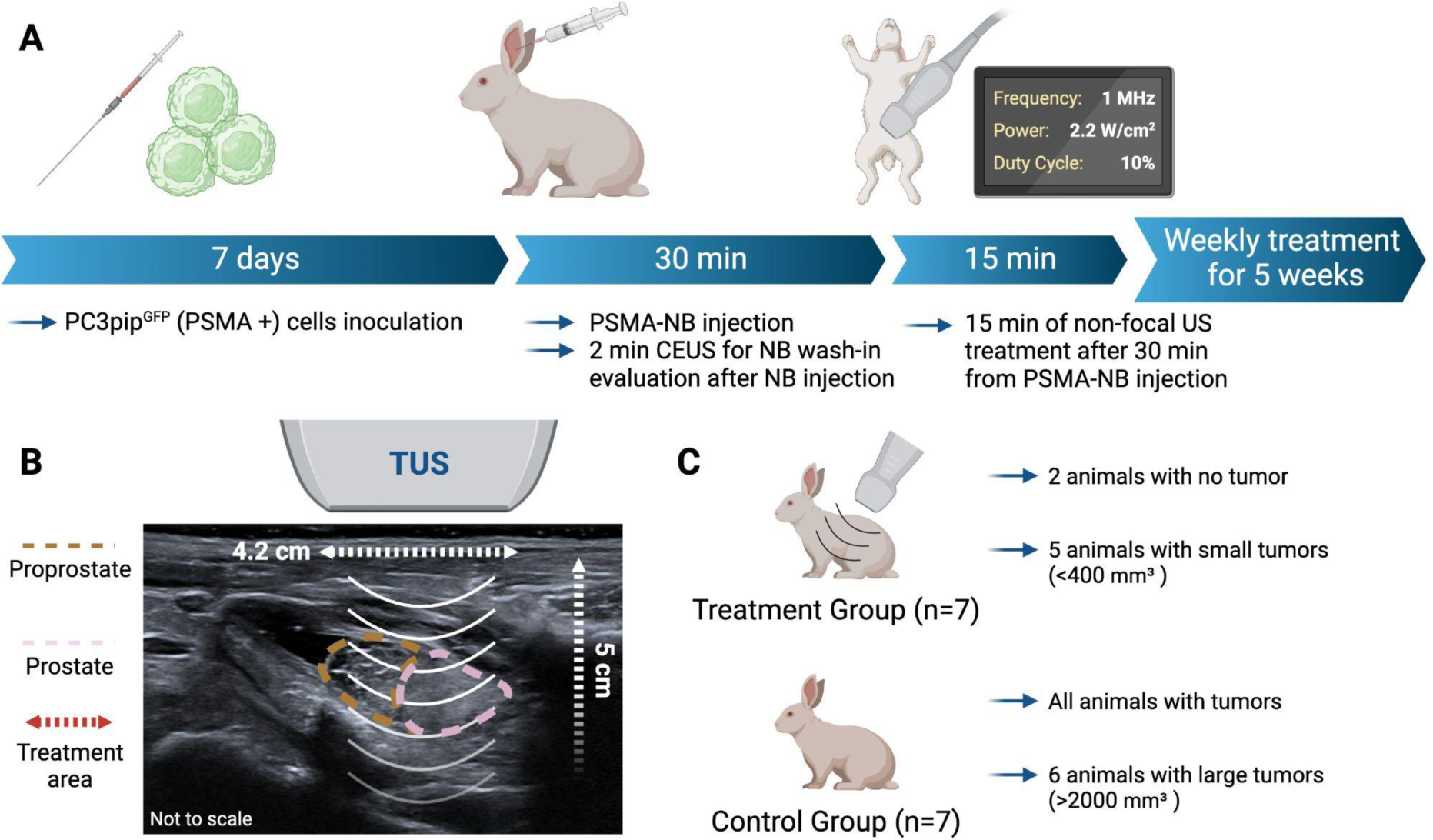
Schematic diagram of the weekly treatment protocol for the rabbits (A). Diagram showing the positioning of the therapeutic ultrasound (TUS) based on the B-mode imaging and illustrating the area of TUS treatment (4.2 cm diameter and 5 cm depth) (B). Summary of the results by group showing that the treatment group presented with less and smaller tumors then the control group (C).

**Figure 6.**
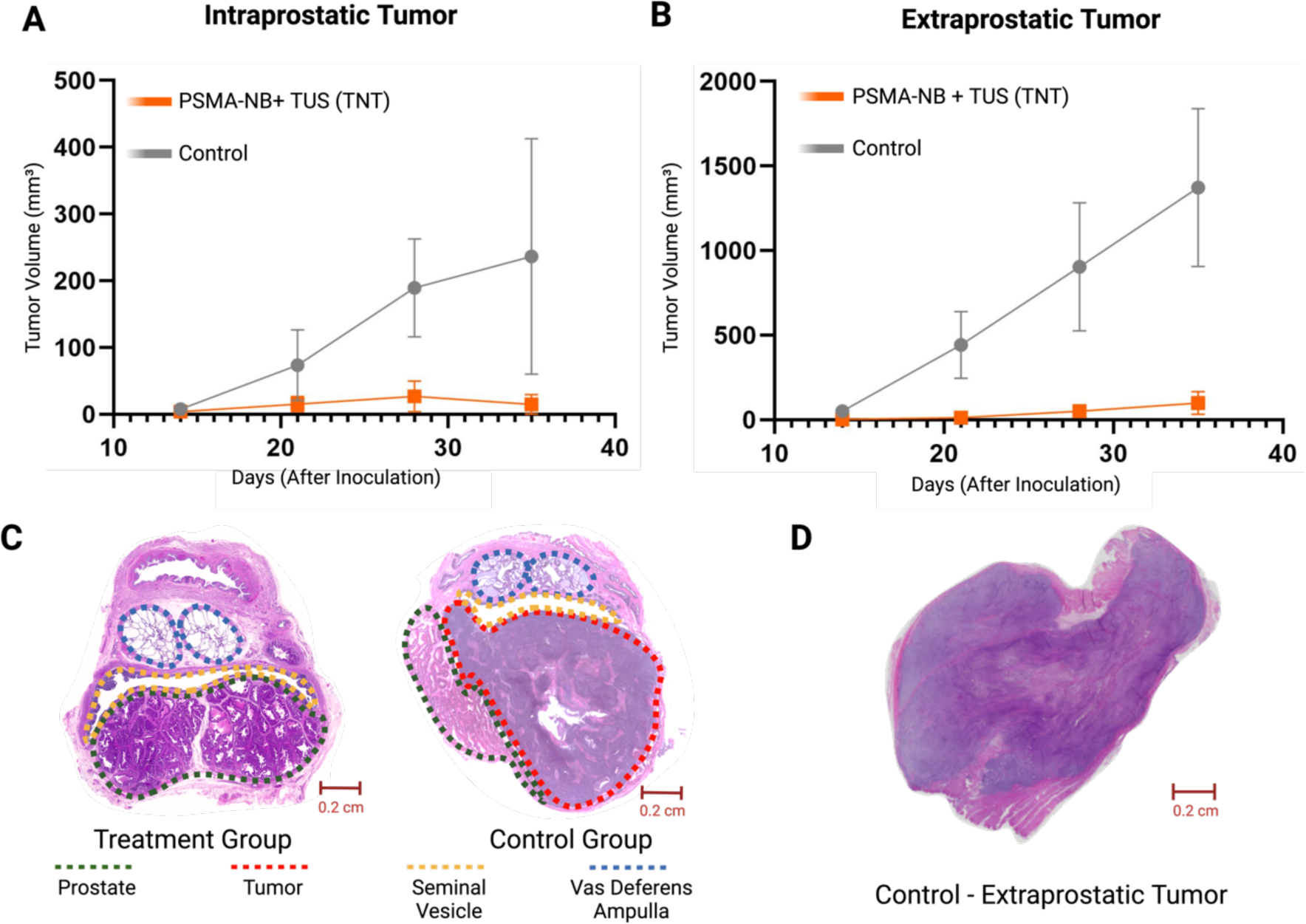
Tumoral growth of intraprostatic tumors (mean ± SEM) (A) and extraprostatic tumors (B) comparing the treatment (PSMA-NB + TUS) and control groups. Representative histology slides (H&E stained) of the prostate gland of the treatment group with no tumor (left) and the control (right) with a large tumor highlighted by the red dashed line (C). Representative histology of an extraprostatic tumor in the control group (PSMA-NB only, no TUS) (D).

## Discussion

In this study, we explored a *drug-free* targeted theranostic approach using PSMA-targeted nanobubbles (PSMA-NBs) in combination with cavitation-induced cell death through unfocused therapeutic ultrasound (TUS) for the treatment of prostate cancer (PCa). Our results demonstrate promising therapeutic efficacy, particularly in inhibiting tumor growth and improving survival rates in both *in vitro* and *in vivo* models, especially in tumors which express PSMA.

Standard treatment for PCa comprises surveillance, localized therapy, and systemic therapy [8,9] each with its specific limitations and side effects. Localized therapies, such as surgery, radiation therapy, and focal therapy, are well-established approaches, but they can lead to significant complications that impact the patient’s quality of life, including issues like morbidity, erectile dysfunction, and incontinence. [6,30–35] Efforts to improve treatment outcomes have led to the development of robot-assisted radical prostatectomy (RARP), which has shown promising results in terms of urinary continence recovery [31]. However, RARP’s widespread implementation is hindered by limited access to robotic surgery platforms globally [32]. Despite these advances, the current treatment options still suffer from suboptimal efficacy and significant side effects, necessitating the exploration of novel therapeutic approaches for PCa.

The combination of treatment modalities in cancer therapy has been explored to minimize side effects while improving treatment efficacy. One approach has been the integration of immunotherapy [35] and photo-thermal therapy (PTT) in preclinical models [36]. Lin et. al [40] reported a novel approach using dual functional nanoparticle and PTT treatment for PCa [37]. While the anti-tumor effect and immune response were enhanced in an *in vivo* subcutaneous mouse tumor model, there is still a lack of information about the side effects. Focal therapies like high-intensity focused ultrasound (HIFU) and laser ablation have been utilized, but they may carry serious side effects [38]. Recently, ultrasound (US) contrast agents, including microbubbles, in combination with US, have been studied to sensitize tumors to radiation therapy, chemotherapy, and gene delivery [39,40]. Ultrasound (US) contrast agents, such as microbubbles, in combination with US have been studied to sensitize tumors to radiation therapy, chemotherapy, and gene delivery [41]. While some studies have shown promising results in inhibiting tumor growth, challenges remain, including off-target effects of the therapeutic of choice due to inevitable systemic distribution, and limitations in efficient tumor cell uptake. In this study we examined a distinct approach leveraging targeted nanobubbles which are internalized into cancer cells via receptor mediated endocytosis. Here, using unfocused low frequency ultrasound, we were able to concentrate the bioeffects of bubble cavitation within individual target cells, which could serve as a complementary, or safer alternative, to extracellular and intravascular cavitation.

To explore intracellular NB cavitation effects, PSMA-NBs were produced with average diameters of 277 nm enabling their uptake into PSMA-expressing cancer cells and aiding in the extravasation into the tumor parenchyma through leaky vasculature [42]. Nanobubble compressibility due to its gas core may also offer additional advantages to extravasation and distribution within the tumor microenvironment. Consistent with our previous findings, confocal imaging revealed a high degree of co-localization of PSMA-NBs in the late endosomal or lysosomal compartments in PSMA+ cells compared to the PSMA-NB in PSMA-cells, confirming their selective uptake via endocytic pathways. Notably, PSMA possesses a unique internalization motif that contributes to a rapid internalization rate of 60% of surface PSMA within 2 hrs [43]. In this study, 1 hr incubation of targeted NBs with PSMA+ cells showed higher cellular uptake compared to the PSMA-cells. The combined treatment of PSMA-NBs with TUS resulted in more perturbed cellular compartments, indicating the destruction of internalized PSMA-NBs and associated damage to cellular compartments following cavitation. Furthermore, PSMA-NBs incubated with PSMA+ cells demonstrated continued localized presence in the cellular compartments of PSMA+ cells up to 24 hrs. These findings underscore the potential of PSMA-NBs as effective carriers for targeted therapies in PCa.

The cell viability effects *in vitro* cell culture were lower than expected resulting in a nominal 25% reduction in viability in PSMA+ cells compared to PSMA-cells after intracellular nanobubble cavitation. The exact reasons for this are unclear, but it is likely that the innate immune system plays an important role in tumor cell death in *in vivo* systems after PSMA-NB and TUS treatment. Further optimization of both NB dose and US energy may be required to optimize in vitro tumor cell death by TNT.

*In vivo* US contrast imaging studies with dual tumor-bearing mice (PSMA+ PC3pip and PSMA-PC3flu tumor) showed higher accumulation of PSMA targeted NBs in the PSMA+ tumor compared to PSMA-tumors after 1 hr. Therefore, the time for the TUS application was selected as 1 hr. to allow targeted NBs to extravasate into the tumor and internalize into tumor cells. Subsequent TUNEL assay results from tumor xenografts demonstrated a high degree of apoptotic cell death with PSMA+ cells treated with combined PSMA-NB and TUS treatment, which indicates selective cell death with combined treatment. Consistent with prior reports [44], the H&E results further provided evidence of the cellular morphology changes in PSMA+ cells with combined treatment. PSMA-tumors had less morphological changes suggesting that the receptor mediated endocytosis plays a role in the TNT mechanism, compared to non-selective accumulation of PSMA-targeted NBs in PSMA-tumor cells

The analysis of long-term tumor growth and the survival analysis indicates that TNT treatment led to decelerated tumor growth in PSMA+ tumor-bearing mice. These effects were dependent on the tumor size at commencement of treatments. In both, PSMA+ and PSMA-groups, the small tumor growth was reduced with the TNT treatment. However, when the TNT treatment was discontinued, the tumors grew rapidly in the PSMA-tumor, indicating recurrence after the treatment, which was not observed in the PSMA+ TNT treatment group. Moreover, Kaplan-Meier survival studies reveal a compelling improvement in the overall survival of PSMA+ tumor-bearing mice after TNT treatment for more than 60 days of post-tumor cell inoculation.

Following evaluation in mice, we assessed the effectiveness of the treatment in an PSMA-expressing orthotopic model of PCa in rabbits [26]. The results demonstrated near-abolished tumor progression in the treatment group especially in extraprostatic lesions. Furthermore, the treatment group also presented with a lower incidence of both intraprostatic and extraprostatic tumors. The effects of intracellular nanobubble cavitation were more prominent in the extraprostatic tumor, we hypothesize this is due to possible better vascularization of the extraprostatic tumors, 0although the precise explanation for this effect is still unclear.

Our previous work with PSMA-NBs (277 nm) demonstrated specific and successful targeting of PSMA-expressing PCa cells both in vitro and in vivo [13,17]. The high degree of co-localization of PSMA-NBs within late endosomal or lysosomal compartments of PSMA+ cells indicated selective uptake via endocytic pathways, yet a lack of significant direct effect on cell viability in vitro. In a similar way, Shen *et al*., conducted research using folate-conjugated NBs (F-NBs) to selectively kill the folate-receptor (FR) positive cervical and lung cancer tumors in a similar approach. However, the NBs used in this study were large (617 nm average diameter) and cationic, which may significantly impact the tumor distribution and cancer cell uptake. In addition, the treatment showed significant cancer cell death in vitro in contrast to the present study, suggesting a different, perhaps complementary mechanism at play [44–46]. Our differential results between lower in vitro cellular killing (24.6 ± 1.4%) and greater in vivo TNT efficacy suggests that there may be an immune-response related mechanism at play, making NBs promising candidates for targeted drug-free therapy specific to prostate cancer.

Despite the promising findings, this study has several limitations. First, the use of athymic mice and immunosuppressed rabbits reduces the potential synergic effect between the nanobubble-based treatments and the immune response, which could have provided valuable insights into the overall mechanism of action and improved overall therapeutic efficacy. Secondly, the choice of a highly aggressive cell line (PC3) for the experiments, although convenient for rapid tumor growth, may not fully represent the typical characteristics of prostate adenocarcinoma. Furthermore, due to resource constraints, the effect of TUS alone or untargeted nanobubbles alone or in conjunction with ultrasound on tumor progression were not evaluated in the rabbit model. This limitation hinders a comprehensive understanding of the specific contribution of PSMA-NBs to the observed therapeutic efficacy. Finally, we have shown strong data demonstrating that tumor tissues that do not express PSMA take up very little PSMA-targeted NB have very little damage. While this implies that normal tissues will also not suffer NB-cavitation damage, it is important to perform studies to determine the extent of impact of NB cavitation on normal cells surrounding tumor cells, which undergo TNT.

## Conclusion

US-mediated destruction of targeted-NBs in mouse PCa xenografts and orthotopic PCa in rabbits shows promise as an effective, highly specific theranostic agent in combination with unfocused TUS cavitation with no extratumoral effects noted on histology. The treatment does not require any additional therapeutic agents and can be used without the need for focused ultrasound instrumentation or magnetic resonance imaging guidance. This increases its safety profile as well as accessibility and decreases complexity of the approach, which could facilitate rapid clinical translation and implementation. Ultimately, the targeted nanobubble therapy strategy has the potential to improve the precision and decrease the side effects of prostate cancer treatment, and can be easily extended to other targets. Further investigation is required to better understand the specific mechanisms of action of intracellular nanobubble cavitation and the role of the immune response in this treatment strategy.

## Abbreviations

ADT: Androgen deprivation therapy
C_3_F_8_: Octafluoropropane
CEUS: Contrast-enhanced ultrasound
CHI: Contrast harmonic imaging
CI: Confidence interval
DBPC: 1,2-dibehenoyl-sn-glycero-3-phosphocholine
DC: Duty cycle
DPPA: 1,2-dipalmitoyl-sn-glycero-3-phosphate
DPPE: 1,2-dipalmitoyl-sn-glycero-3-phosphoethanolamine
DSPE-mPEG: 1,2-distearoyl-snglycero-3-phosphoethanolamine-N-[methoxy(polyethylene glycol)-2000]
F-NB: Folate-conjugated nanobubble
FR: Folate receptor
GFP: Green-fluorescent protein
H&E: Hematoxylin and eosin
HIFU: High-Intensity Focused Ultrasound
HPLC: High-performance liquid chromatography
IACUC: Institutional Animal Care and Use Committee
MALDI-TOF-MS: Matrix-assisted laser desorption/ionization time-of-flight mass spectroscopy
MRI: Magnetic resonance image
NB: Nanobubble
OCT: Optimal-cutting-temperature compound
PBS: Phosphate-buffered saline
PCa: Prostate cancer
PDT: Photodynamic therapy
PSMA: Prostate-Specific Membrane Antigen
PSMA-NB: Prostate-Specific Membrane Antigen Targeted Nanobubble
PTT: Photo-thermal therapy
RARP: Robot-assisted prostatectomy
RMM: Resonant mass measurement
RNA: Ribonucleic acid
ROI: Region-of-interest
RPMI1640: Roswell Park Memorial Institute 1640 Medium
RR: Relative risk
SBRT: Stereotactic body radiotherapy
SD: Standard deviation
SE: Standard error of the mean
TNT: Targeted nanobubble therapy
TRUS: Transrectal ultrasound
TUNEL: Terminal deoxynucleotidyl transferase dUTP nick end labeling
TUS: Non-focal therapeutic ultrasound
UMND: US-mediated NB destruction
US: Ultrasound

## Acknowledgements

We are grateful to Kristie Brock, Melissa Dusek, Melissa Carr, Shelly Weisman, Leonardo K. Bittencourt and Felipe Matsunaga for their support and assistance with the rabbit studies. Dana Wegierak’s help with the manuscript review is also highly appreciated. Felipe Berg is supported by the Marcos Lottenberg & Marcus Wolosker International Fellowship for Physicians Scientist - Case Western Reserve University. We acknowledge the support from Siemens Healthineers for providing the ultrasound device for the rabbit studies and the funding source of this study: National Institute of Biomedical Imaging and Bioengineering (NIBIB) under Award No. 5R01-EB025741 (A.A.E. and J.P.B.) and the Case-Coulter Translational Research Partnership/Wallace H. Coulter Foundation (A.A.E. and J.P.B.). Figures were made with BioRender.com.

