## Supplemental Material for "Ultrasound-mediated drug-free theranostics for treatment of prostate cancer"

[1]Department of Radiology, Case Western Reserve University, Cleveland, OH, United States [2]Instituto Israelita de Ensino e Pesquisa, Hospital Israelita Albert Einstein, São Paulo, SP, Brazil [3]Department of Biomedical Engineering, Case Western Reserve University, Cleveland, OH, USA

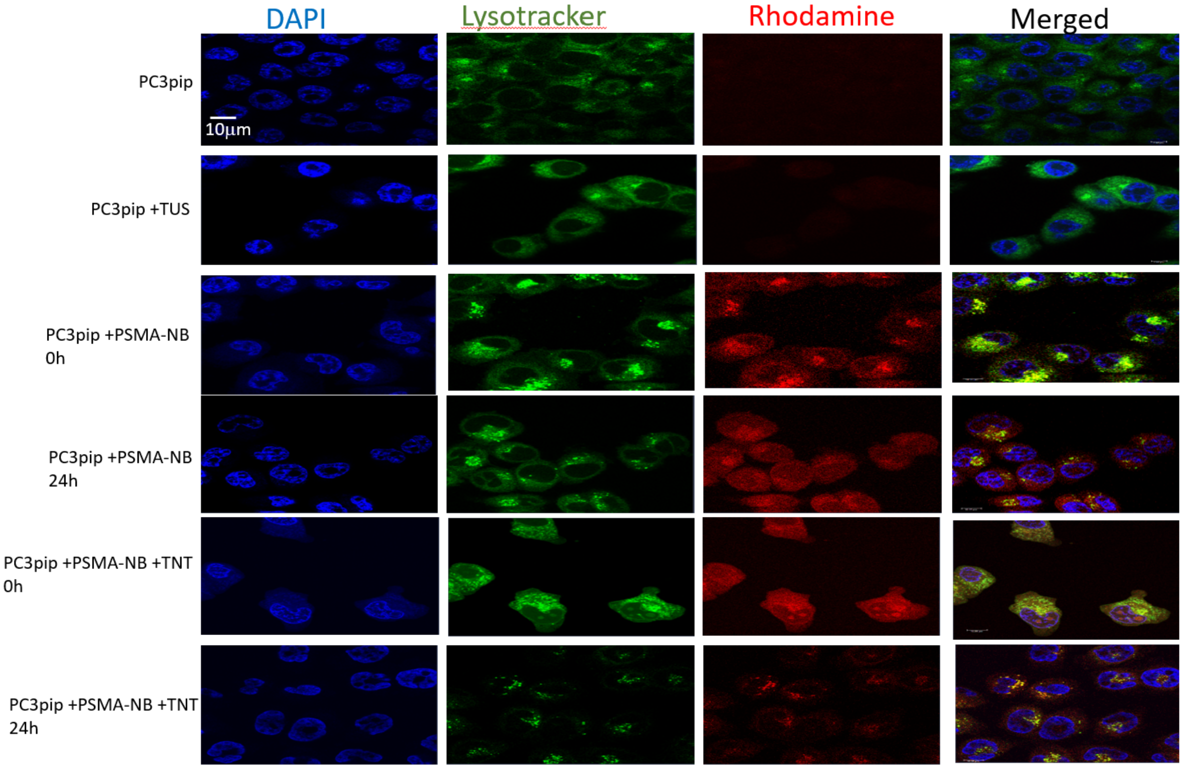


Figure S1. Confocal images of PSMA-NB distribution in PSMA+ PC3pip cells after 1 hr co-incubation with PSMA-NB (t=0), 24h post incubation, and after co-incubation with PSMA-NB and the therapeutic US applied (for 10min, 2.2 power, 3MHz, and 10% duty cycle); t=0 and t=24 post-treatment. 100X (blue-nuclei, red-NB, green-late endosome/lysosomes, and yellow-co-localized NBs with endosome/lysosomes).

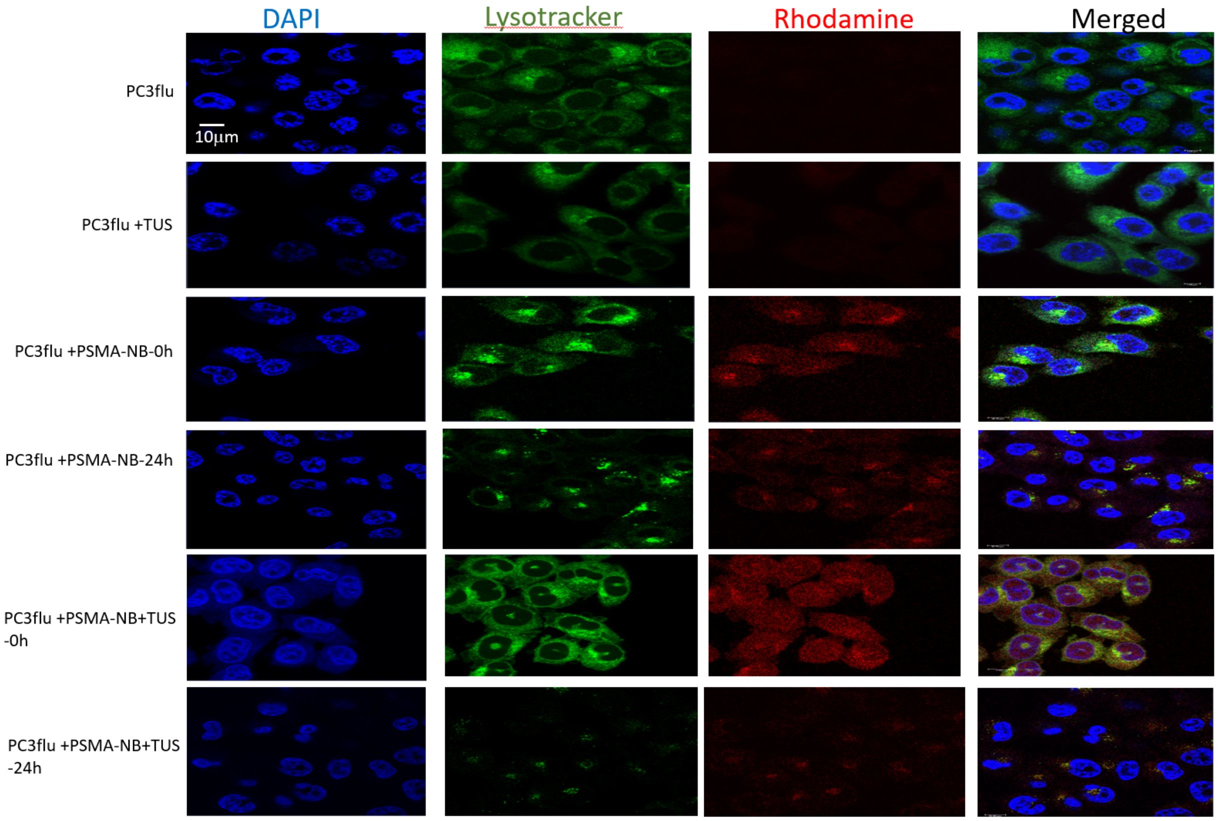


Figure S2. Confocal images of PSMA-NB distribution in PSMA- PC3flu cells after 1 hr co-incubation with PSMA-NB (t=0), 24h post incubation, and after co-incubation with PSMA-NB and the therapeutic US applied (for 10min, 2.2 power, 3MHz, and 10% duty cycle); t=0 and t=24 hrs post-treatment. 100X (blue-nuclei, red-NB, green-late endosome/lysosomes, and yellow-co-localized NBs with endosome/lysosomes).

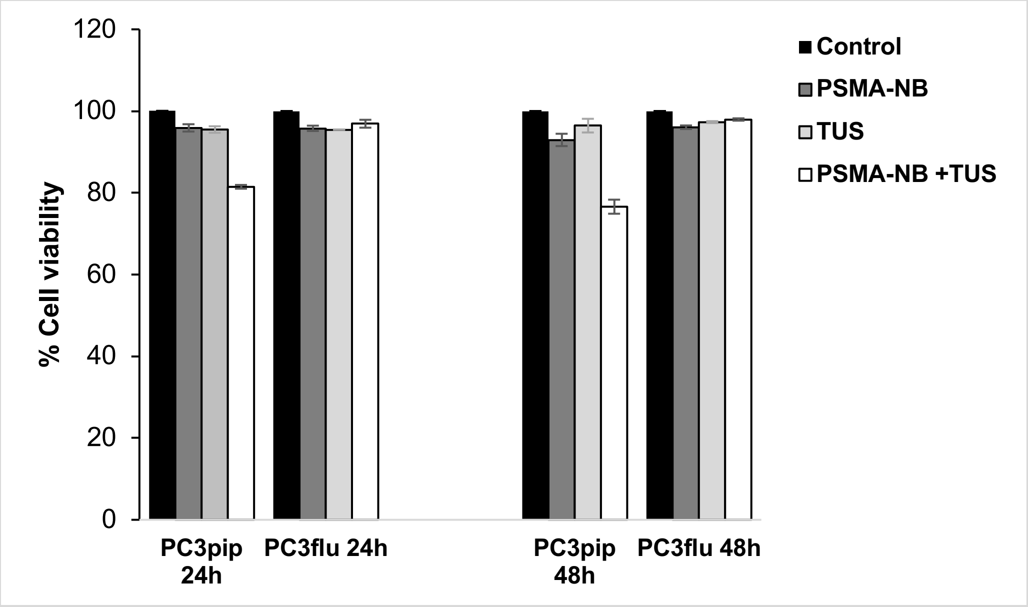


Figure S3. The cell viability measured using WST-1 assay. PSMA+ PC3pip cells treated with both PSMA-NB and TUS showed more cell death at 24 hrs and 48hrs compared to PSMA- PC3flu cells.

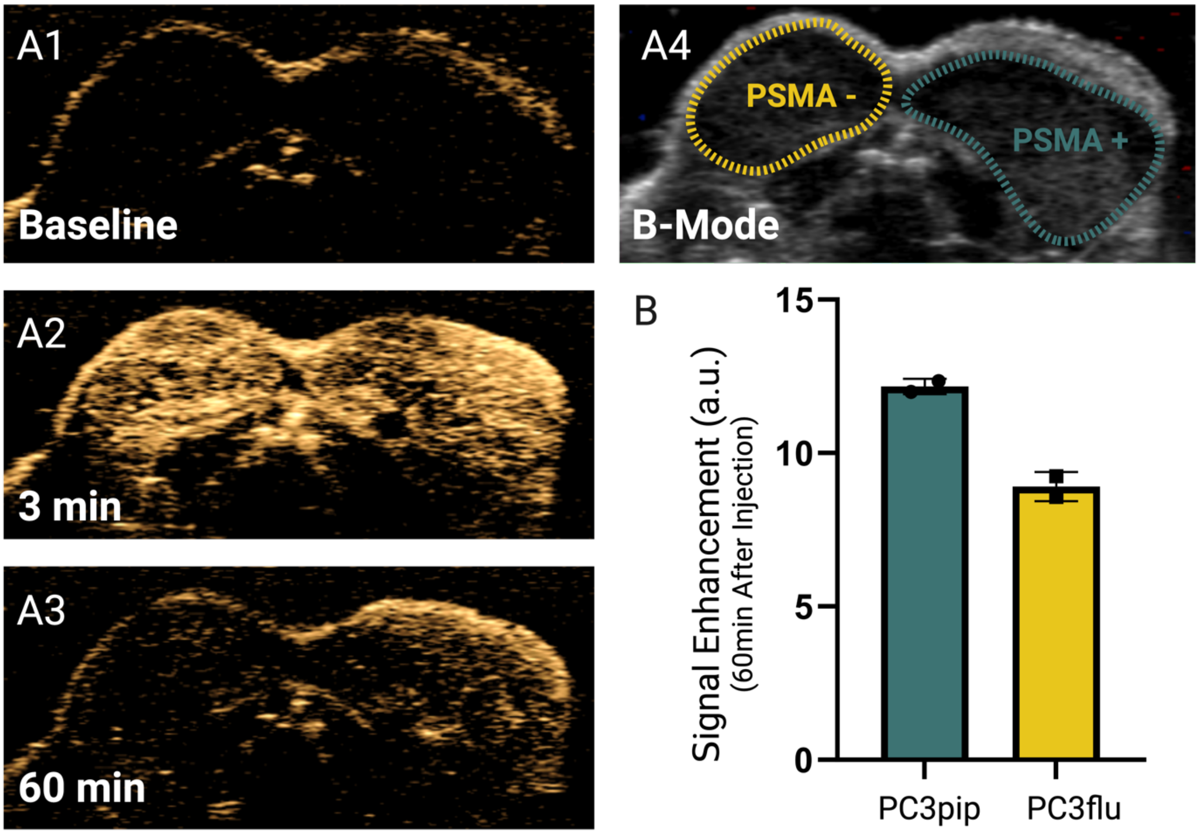


Figure S4. US images of PSMA-NB in PSMA+ and PSMA- tumors showing the contrast enhancement before the injection (A1), 3 min after injection (A2) and 1h after injection (A3). Quantified signal enhancement 1h post-injection (B).

(a).

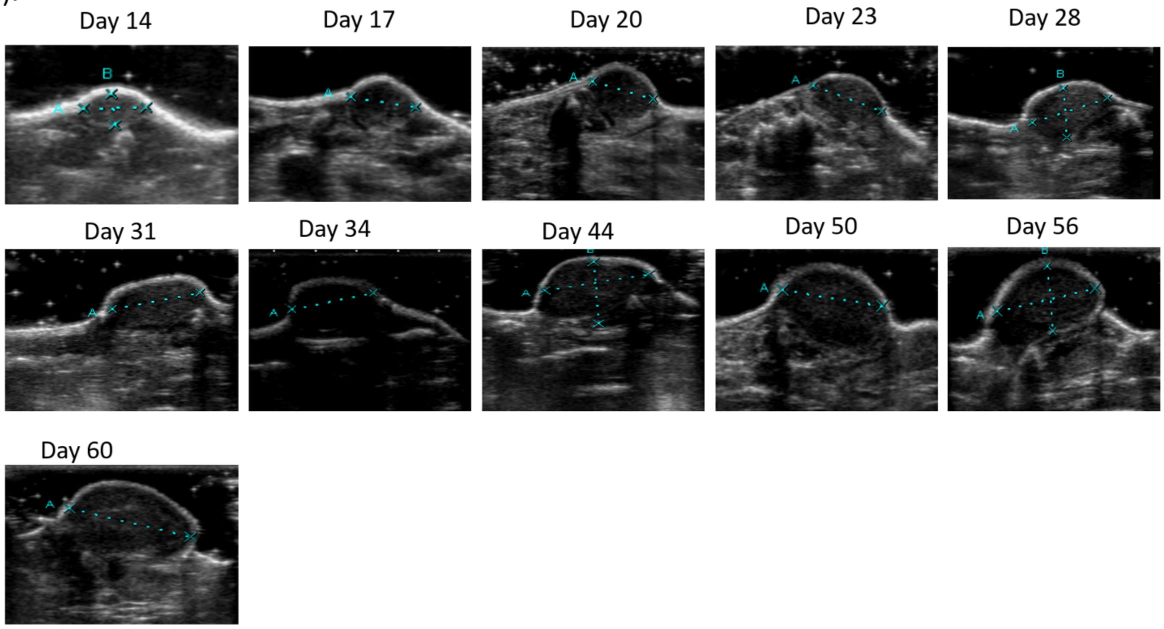
Day 14
Day 17
Day 20
Day 23
Day 28
Day 31
Day 34
Day 44
Day 50
Day 56
Day 60

(b).

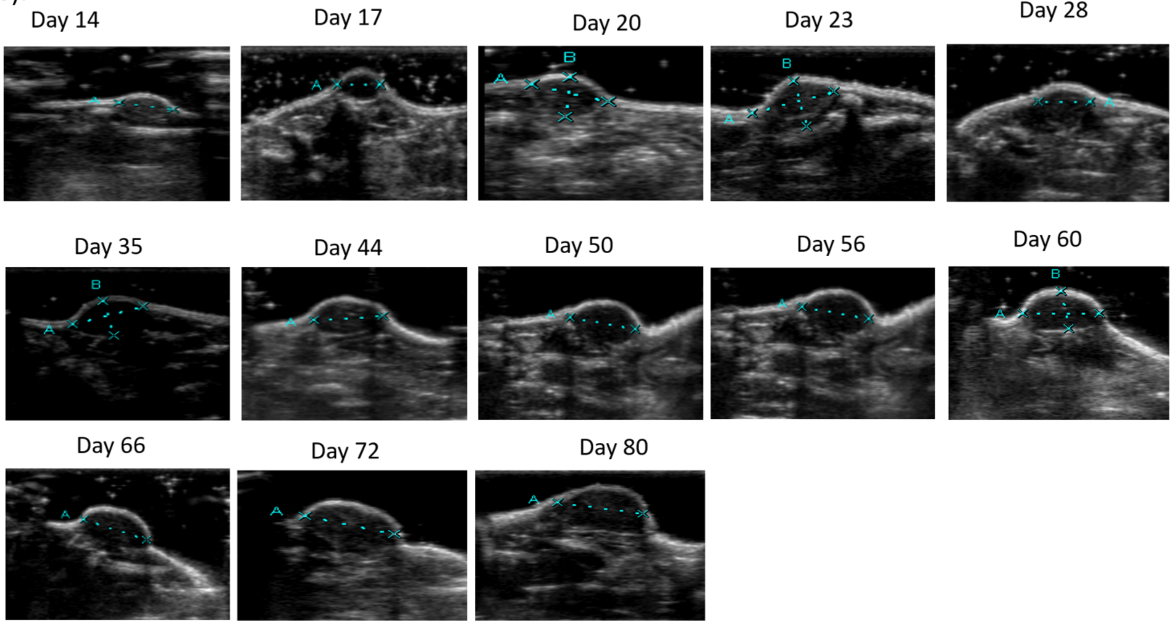
Day 14
Day 17
Day 20
Day 23
Day 28
Day 35
Day 44
Day 50
Day 56
Day 60
Day 66
Day 72
Day 80

(c).

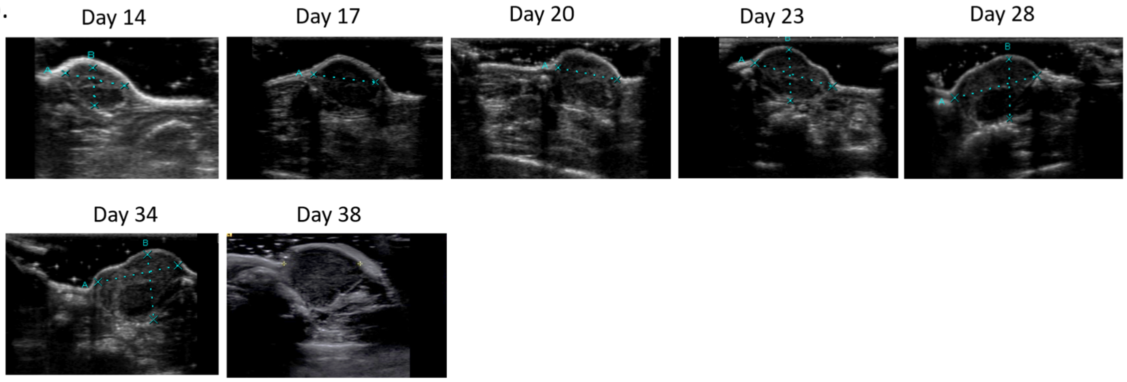


(d).

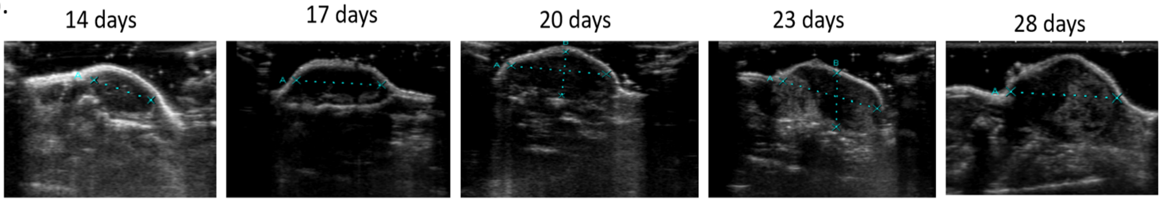


Figure S5. (a) Representative B-mode images of PSMA+ PC3pip tumor treated with PSMA-NB combined with therapeutic US (TNT) as a function of time after the tumor inoculation. On days 14, 17, 20, and 23 the treatment was applied. (b) B-mode images of PC3pip tumor treated with PSMA-NB combined with therapeutic US (TNT) as a function of time after the tumor inoculation, which demonstrated the retardation of tumor growth with time with increased survival date (80 days). On days 14, 17, 20, and 23 the treatment was applied. (c) Representative B-mode images of PSMA+ PC3pip tumor treated with therapeutic US (TUS) as a function of time after the tumor inoculation. On days 14, 17, 20, and 23 the treatment was applied. (d) Representative B-mode images of untreated PC3pip tumor as a function of time after the tumor inoculation. on days 14, 17, 20, and 23 the PBS was only injected.

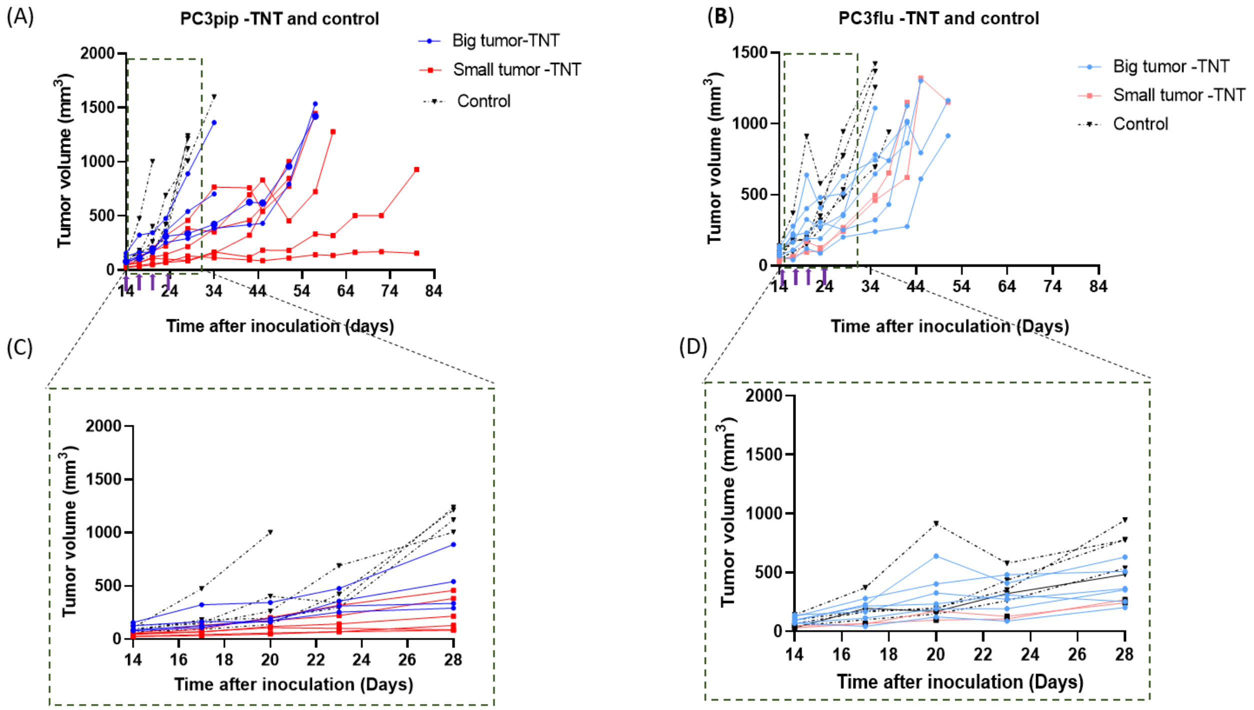


Figure S6. Tumor growth curve of PSMA+ PC3pip and PSMA- PC3flu tumor mice treated with TNT (PSMA-NB + TUS) treatment and control group (a) PC3pip tumor mice with big tumors (> 60mm$^3$ initial tumor volume) and small tumors (< 60mm$^3$ initial tumor volume) treated with TNT treatment and control group (b) PC3flu tumor mice with big tumors (> 60mm$^3$ initial tumor volume) and small tumors (< 60mm$^3$ initial tumor volume) treated with TNT treatment and control group. Zoomed to show the tumor growth for 28 days for (c) PSMA+ PC3pip tumor and (d) PSMA- PC3flu tumor.

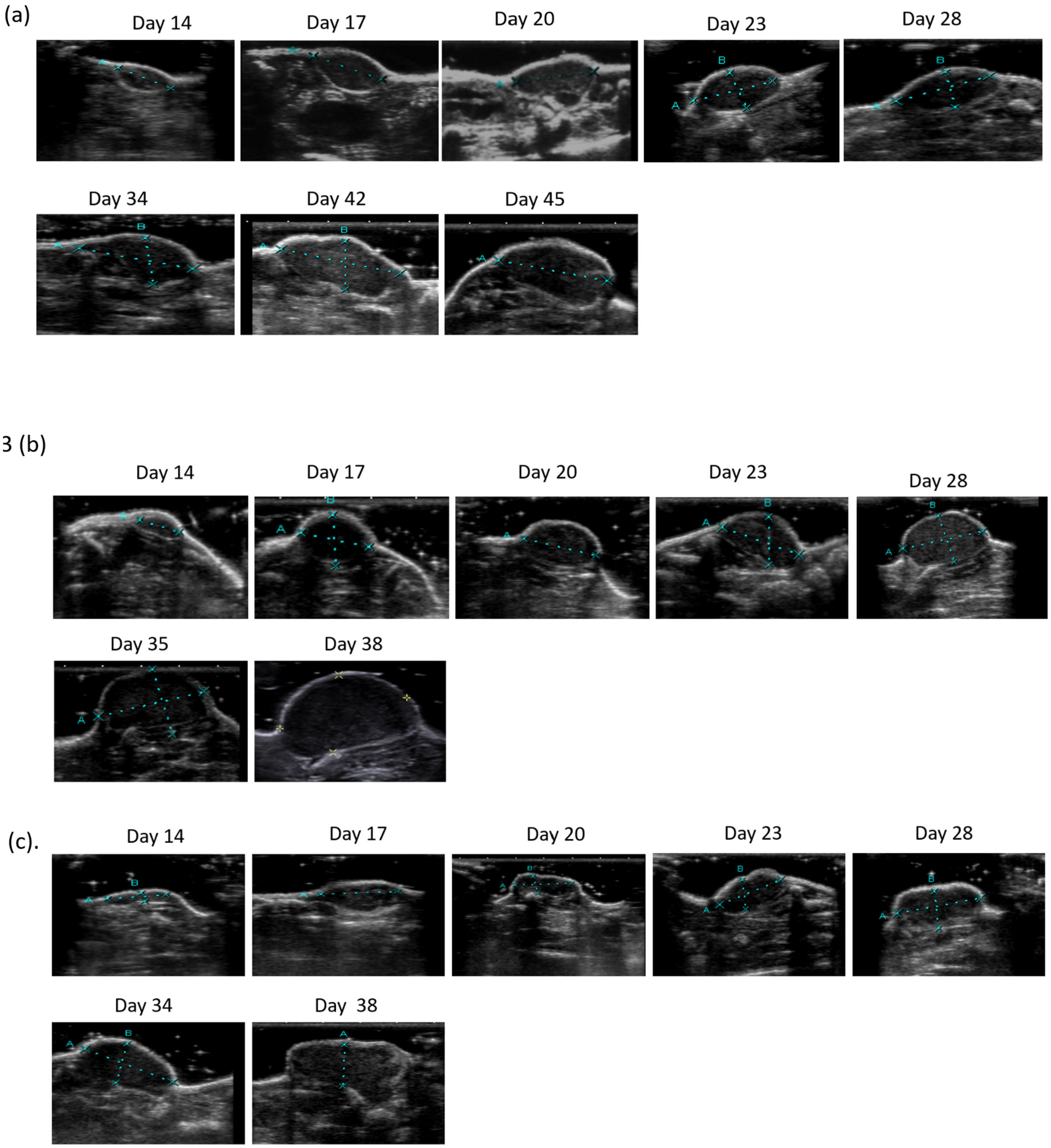


Figure S7. (a) Representative B-mode images of PSMA- PC3flu tumor treated with PSMA-NB combined with Therapeutic US (TNT) as a function of time after the tumor inoculation. On days 14, 17, 20, and 23 the treatment was applied. (b) Representative B-mode images of PSMA- PC3flu tumor treated with Therapeutic US (TUS) as a function of time after the tumor inoculation. On days 14, 17, 20, and 23 the treatment was applied. (c) Representative B-mode images of untreated PC3flu tumor as a function of time after the tumor inoculation. On days 14, 17, 20, and 23 the PBS was only injected.

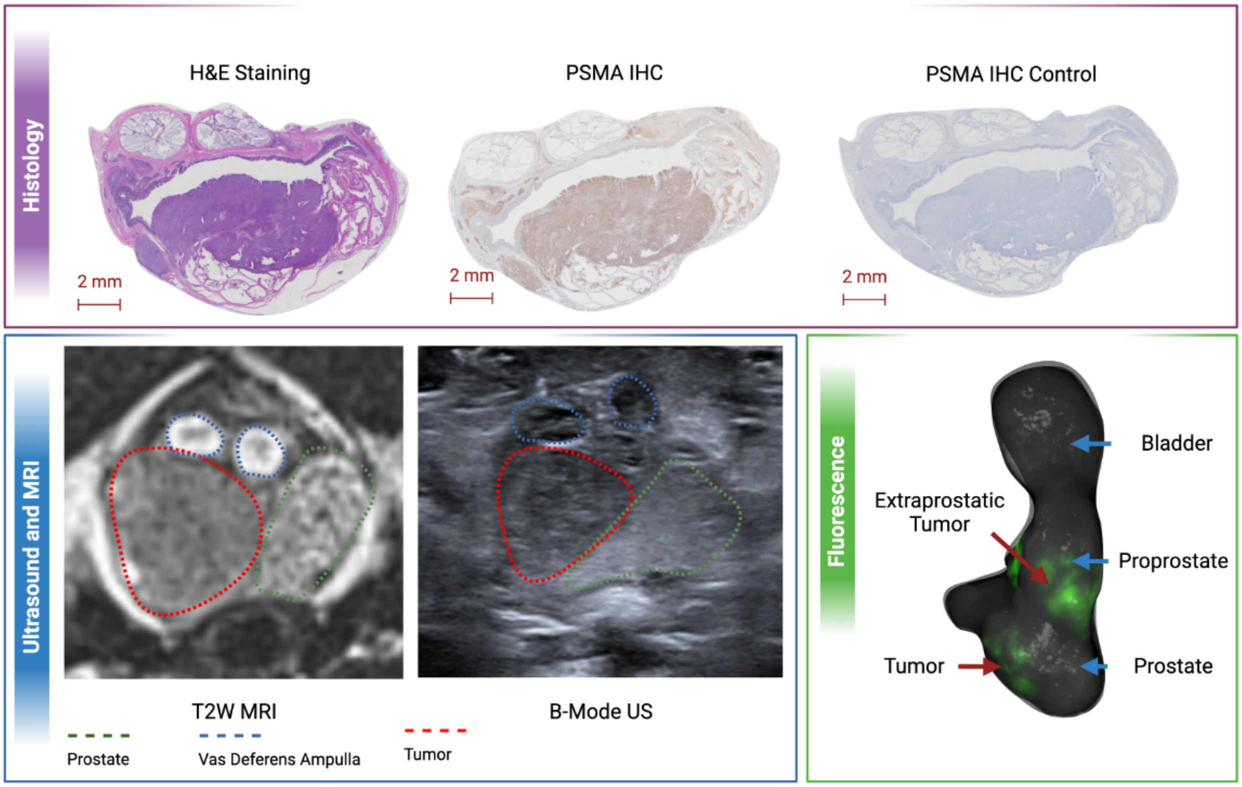


Figure S8. Histology (top) with H&E, PSMA IHC for the tumors. Ultrasound and MRI of the tumors for lesion localization (bottom left). Fluorescence (green fluorescent protein) image for tumor localization on the harvested gland and tissues.

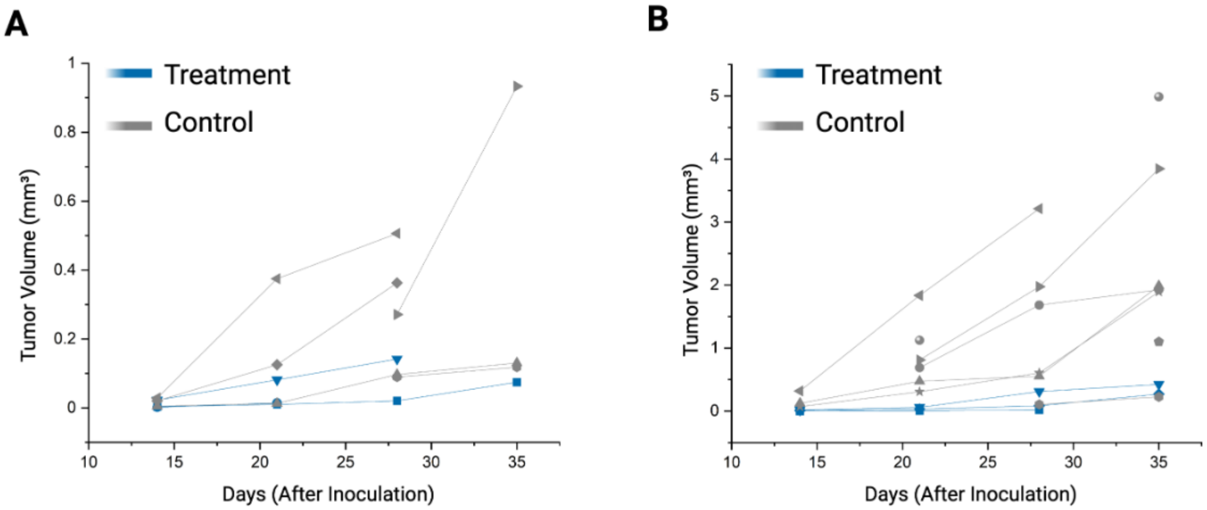


Figure S9. Tumoral individual growth curves (per tumor) for the intraprostatic lesions (A) and extraprostatic lesions (B).
